# Notch3 regulates pericyte phenotypic plasticity in colorectal cancer

**DOI:** 10.1101/2025.06.04.657823

**Authors:** Niki Chalkidi, Athanasia Stavropoulou, Vasiliki-Zoi Arvaniti, Christina Paraskeva, Artemis Monogyiou, Maria Sakkou, Christoforos Nikolaou, Vasiliki Koliaraki

## Abstract

Pericytes undergo phenotypic alterations that influence cancer progression, yet the molecular mechanisms governing these changes remain poorly understood. Here, we investigated the role of Notch3 signaling in pericyte phenotype and functions in colorectal cancer (CRC). Using lineage tracing approaches, we showed that tumor pericytes originate from normal tissue-resident pericytes, which proliferate inside tumors. In vivo genetic manipulation revealed that Notch3 pathway activation promotes pericyte proliferation, while suppressing contractile protein expression, and leads to increased endothelial cell proliferation. In contrast, Notch3 deletion leads to reduced endothelial proliferation and a significant decrease in. This effect is associated with a shift toward a contractile phenotype. Single-cell RNA sequencing analysis uncovered significant pericyte heterogeneity in both mouse colitis-associated cancer and human CRC. It specifically identified functionally distinct subpopulations characterized by differential Notch3 activity, which supported our in vivo and in vitro findings. Our results establish Notch3 as a key regulator of pericyte phenotypic plasticity in CRC and suggest that targeting this pathway could represent a promising strategy for improving therapeutic outcomes through vascular normalization.

## Introduction

Colorectal cancer (CRC) remains a significant global health burden, ranking as the third most diagnosed cancer and the second leading cause of cancer-related mortality worldwide (*1, 2*). This highlights the need for novel therapeutic approaches based on deeper understanding of CRC biology. Recent research has uncovered the crucial role of the tumor microenvironment (TME) in CRC progression, metastasis, and therapy resistance (*3, 4*). The TME consists of diverse cell populations, including fibroblasts, immune cells, endothelial cells, and pericytes, which collectively influence tumor behavior through complex intercellular communications and extracellular matrix remodeling (*4*).

Pericytes are specialized perivascular cells embedded within the basement membrane of microvessels, where they establish direct physical contact with endothelial cells (*5*). In physiological conditions, pericytes play essential roles in vascular development, stabilization, and maturation by regulating endothelial cell proliferation, vessel diameter, and basement membrane deposition. The recruitment and differentiation of pericytes along nascent vessels are tightly regulated by several signaling pathways, such as PDGF-B, TGFβ, and angiopoietin. Disruption of these pathways leads to pericyte deficiency, vascular instability, and subsequent pathologies, emphasizing the importance of proper pericyte function in vascular homeostasis (*5*). In cancer, pericytes undergo significant alterations in their phenotype, distribution, and functionality, contributing to tumor angiogenesis by regulating vascular integrity, permeability, and remodeling (*6*). Tumor vessels typically exhibit abnormal pericyte coverage characterized by loose attachment to endothelial cells, cytoplasmic processes extending into the tumor parenchyma, and altered expression of marker proteins. These abnormalities contribute to vessel leakiness, increased interstitial fluid pressure, and hypoxia, ultimately promoting tumor progression and limiting drug delivery (*7*). In addition, pericyte phenotype switching in tumors has emerged as an important process influencing cancer progression. Upon exposure to tumor-derived factors, pericytes may acquire a more proliferative state, transdifferentiate into fibroblast-like cells, or adopt contractile properties resembling smooth muscle cells (*8–10*). The molecular mechanisms governing these phenotypic transitions remain incompletely understood, particularly in the context of CRC. Furthermore, the precise origin of tumor pericytes in CRC has been a subject of debate, with potential sources including resident pericytes, recruited bone marrow-derived pericyte precursors or transdifferentiation from other stromal cell types, mainly endothelial cells (*11–13*). Understanding these processes is crucial for developing strategies to normalize the tumor vasculature and improve therapeutic outcomes (*14, 15*).

The Notch signaling pathway represents a highly conserved cell-cell communication mechanism involved in various aspects of development, tissue homeostasis, and disease, including cancer (*16–18*). In mammals, the pathway consists of four transmembrane receptors (Notch1-4) and five ligands (Jagged1-2, Delta-like 1, 3, and 4). Activation of Notch receptors through interaction with ligands on adjacent cells triggers a series of proteolytic cleavages, resulting in the release of the Notch intracellular domain (NICD). The NICD subsequently translocates to the nucleus, where it forms a transcriptional complex with the DNA-binding protein RBPJ and co-activators to regulate target gene expression (*19*). Among Notch receptors, Notch3 has gained particular attention for its role in vascular development and homeostasis. Notch3 is predominantly expressed in vascular smooth muscle cells (vSMCs) and pericytes, where it regulates cell differentiation, maturation, and survival (*20*). Mutations in the *NOTCH3* gene cause cerebral autosomal dominant arteriopathy with subcortical infarcts and leukoencephalopathy (CADASIL), a hereditary stroke disorder characterized by progressive degeneration of vascular smooth muscle cells (*21*). In mice, deletion of Notch3 leads to abnormal maturation of vSMCs and reduced mural cell recruitment and maturation in the retinal vasculature (*20, 22*). These findings highlight the importance of Notch3 signaling in maintaining the structural and functional integrity of the vasculature. In cancer, Notch3 signaling exhibits context-dependent effects, functioning either as an oncogene or a tumor suppressor depending on the tissue type and cellular context (*23*). Several studies have reported Notch3 overexpression in various malignancies, including ovarian, breast, and lung cancers, where it promotes tumor cell proliferation, survival, and stemness (*23, 24*). In colorectal cancer, elevated Notch3 expression correlates with advanced tumor stage and poor prognosis (*23, 25, 26*). Notch3 upregulation specifically in cancer cells results in increased tumour growth, while its inhibition leads to reduced proliferation and invasiveness both in xenografts and a mouse model of CRC (*26–28*). However, the specific role of Notch3 in regulating pericyte behavior within the CRC microenvironment remains largely unexplored, representing a significant knowledge gap in the field.

In this study, we employed single-cell RNA sequencing, lineage tracing, and functional assays to investigate the role of pericytes in CRC development using the AOM/DSS colitis-associated cancer model. We identified a significant enrichment of pericytes in the tumor microenvironment compared to normal colon tissue and showed that Notch3 signaling plays an important role in regulating pericyte proliferation, phenotype, and interaction with endothelial cells. Our findings provide novel insights into the contribution of pericytes to CRC progression and suggest that targeting Notch3 signaling in pericytes may represent a promising therapeutic approach for advanced colorectal cancer.

## Results

### Single cell analysis reveals enrichment of pericytes in the microenvironment of colorectal cancer

In the AOM/DSS model of colitis-associated cancer (CAC) model, carcinogenesis is initiated by a single injection of azoxymethane (AOM), which induces random mutations. This process is further enhanced by cycles of inflammation and regeneration triggered by cyclic administration of dextran sodium sulfate (DSS) in drinking water. The resulting tumors are predominantly adenomas located in the middle and distal colon (*29*). Having previously used genetic models to delineate the role of fibroblast-specific pathways in this process, we employed here a high-throughput single-cell approach to analyze the stromal component of these tumors. To this end, we isolated non-epithelial, non-immune (EpCAM-CD45-) cells from AOM/DSS colonic adenomas (tumors from 11 male and female mice) and normal healthy colon tissue (pooled from one male and one female mouse).

Isolated stromal cells were used for library preparation on the 10x Genomics platform and subsequent single cell RNA sequencing. After excluding glial cells (*Gfap+S100b*+) and interstitial cells of Cajal (*Kit*+*Ano1*+), and any remaining immune (*Ptprc*+) and epithelial/cancer (*Epcam*+) cells, we integrated the data across the two conditions using Harmony (*30*). 12.692 (normal: 6.018, AOM/DSS: 6.674) stromal cells were included in the subsequent bioinformatic analysis, after quality control and doublet exclusion. Clustering and UMAP visualization revealed 6 broad clusters, which correspond to known cell types, including blood and lymphatic endothelial cells, pericytes, smooth muscle cells (SMCs), fibroblasts, and a distinct proliferating stromal cluster (Figure 1A-C and Supplementary Data 1).

**Figure 1.**
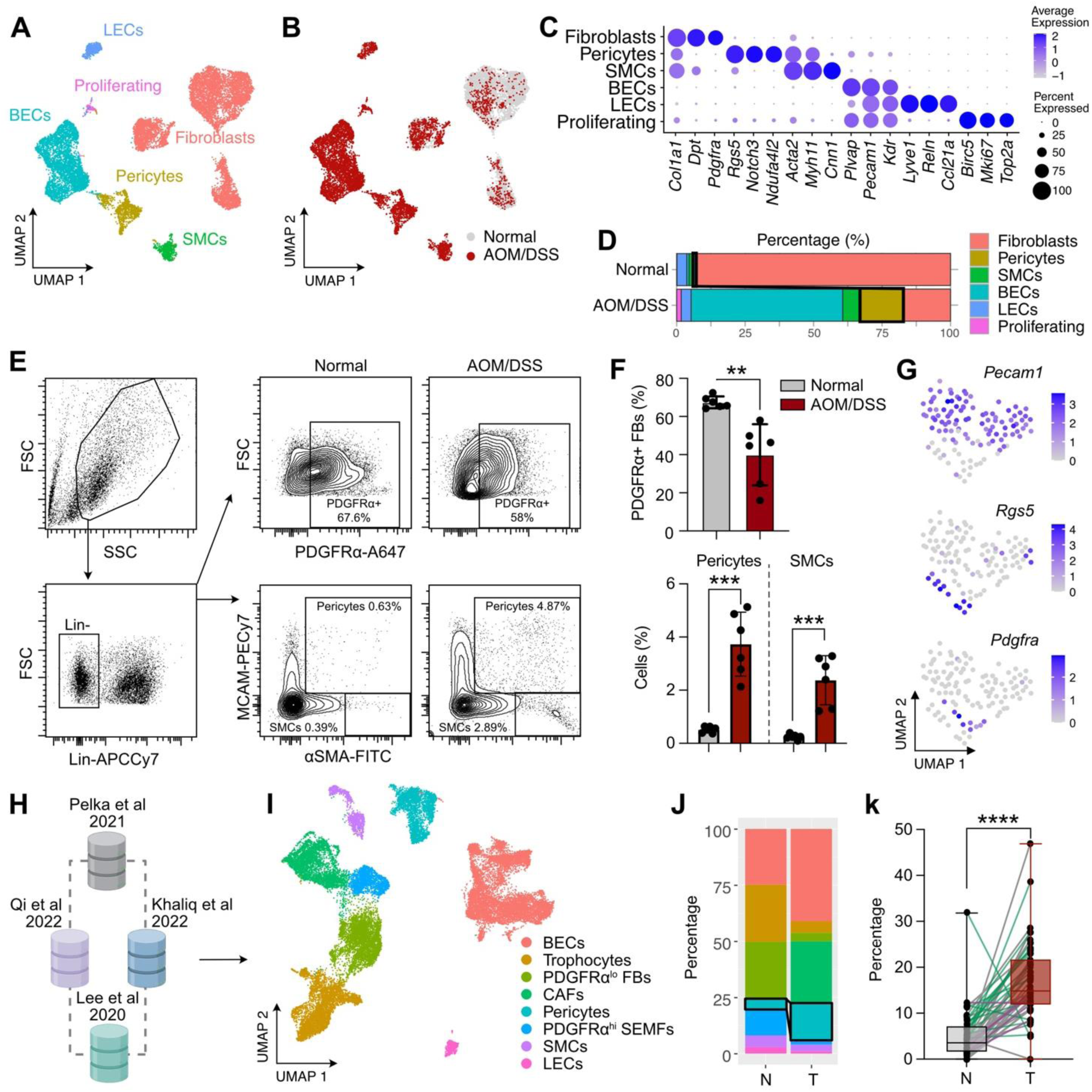
Pericytes are enriched in murine colitis-associated cancer and human CRC. UMAP visualization of stromal cells in AOM/DSS tumors and normal colon, based on (A) cluster identity and (B) sample of origin (Normal or AOM/DSS). (C) Dot plot showing the expression of marker genes for the different stromal clusters (p.adj ≤ 0.05). (D) Percentage of each stromal cluster per sample. (E) Representative FACS gating strategy, plots and (F) quantifications of PDGFRα^+^ fibroblasts, MCAM^+^αSMA^+^ pericytes, and αSMA^hi^ smooth muscle cells (SMCs) in the normal mouse colon and AOM/DSS tumors (n = 6, one-way ANOVA). (G) Feature plots showing marker gene expression in subclustered proliferating stromal cells. (H) Schematic showing the four single-cell RNA sequencing studies of human colorectal cancer. (I) UMAP visualization of stromal cells in the integrated human CRC dataset. Cluster identification was performed in accordance with the mouse data using a similar set of marker genes. (J) Relative abundance of pericytes in the integrated human CRC stroma. (K) Percentage of pericytes in tumor (T) samples and adjacent normal (N) mucosa. Only samples that included both states were used. Individual patient samples with <10 cells were excluded from the analysis (n = 49 pairs, ****p<0.0001).

Blood endothelial cells could be further subdivided in capillary (*Rgcc*+ CapECs), venous (*Ackr1*+ VenECs), and arterial (*Sema3g*+ ArtECs) endothelial cells, consistent to previous studies (Supplementary Figure 1A-C and Supplementary Data 1) (*31*). Similarly, fibroblasts were subdivided into three main clusters: PDGFRα^hi^ subepithelial myofibroblasts (SEMFs), PDGFRα^lo^CD81-fibroblasts and PDGFRα^lo^CD81+ trophocytes (Supplementary Figure 2A-D and Supplementary Data 1) (*32, 33*).

Notably, all identified clusters contained cells from both normal tissue and tumors, albeit at different proportions. Tumor samples showed increased numbers of blood endothelial cells, pericytes, and smooth muscle cells, with a corresponding decrease in fibroblasts (Figure 1D). This observation was further verified by FACS analysis, which confirmed an increase in MCAM+αSMA+ pericytes and αSMA^hi^ SMCs, along with a decrease in PDGFRα+ fibroblasts (Figure 1E-F). An increase in MCAM+αSMA+ cells in the AOM/DSS model has also been previously reported (*34*). In addition, tumor samples had more proliferating stromal cells, primarily consisting of endothelial cells and pericytes in line with their observed increase (Figure 1G). Single-cell transcriptomic data integration of human CRC stromal cells from four published studies showed similar changes in stromal cell proportions, and in particular the significant increase in pericytes (*35–38*) (Figure 1H-K, Supplementary Figure 3, and Supplementary Data 1). These findings show that intestinal carcinogenesis is accompanied by a significant increase in cancer-associated pericytes in both mice and humans.

### Lineage tracing confirms pericyte expansion in colitis-associated cancer

Given the increased abundance of cancer-associated pericytes and the presence of proliferating pericytes, we next investigated whether normal pericytes are a source of tumor pericytes and whether they proliferate within tumors. To address these questions, we utilized the PDGFRβ-CreERT2 mouse (*39*) in combination with the mTmG reporter strain (*40*) and performed lineage tracing analysis (Figure 2A). Tamoxifen was administered for five consecutive days prior to the initiation of the AOM/DSS protocol. At the end of the protocol, we analyzed both normal colon tissue and tumors by flow cytometry and confocal imaging. Our analysis revealed a significant approximately 3-fold increase in labeled PDGFRβ+MCAM+ pericytes within tumors compared to the normal tissue (Figure 2B-D). However, we noted that the PDGFRβ-CreERT2 strain also targets intestinal fibroblasts, as previously described (*41*).

**Figure 2.**
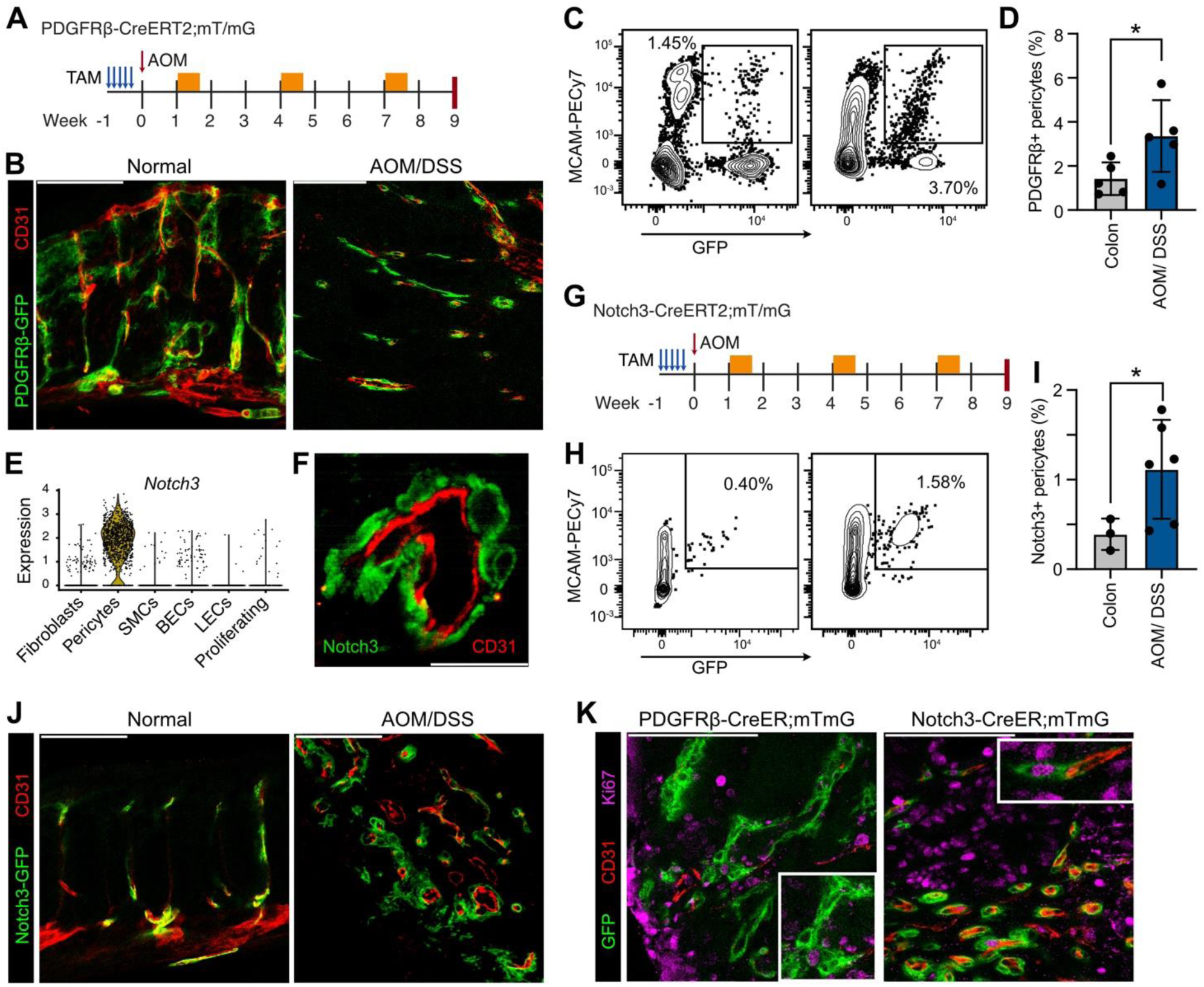
Fate mapping of colonic pericytes during colitis associated carcinogenesis. (A) Schematic representation of experimental setup for PDGFRβ+ lineage tracing (Orange boxes represent DSS administration, arrows represent injections, and the red line the end of the protocol). (B) Representative confocal images of GFP+ cells in the normal colon and AOM/DSS tumors (n = 5) (Scale bar: 100 μm). CD31 staining was used to label endothelial cells. (C) Representative FACS plots and (D) quantification of PDGFRβ-GFP+ MCAM+ pericytes in Lin-cells (n = 5, unpaired two-way t-test). (E) Violin plot showing *Notch3* expression in our single cell dataset. (F) Representative confocal image showing Notch3 staining around CD31+ blood vessels (Scale bar: 20 μm). (G) Schematic representation of experimental setup for Notch3+ lineage tracing. (H) Representative FACS plots and (I) quantification of Notch3-GFP+ MCAM+ pericytes in Lin-cells (n = 3-6, Welch’s t-test). (J) Representative confocal images of Notch3-GFP+ cells in the normal colon and AOM/DSS tumors (n = 6) (Scale bar: 100 μm). (K) Representative confocal images of PDGFRβ-GFP+ and Notch3-GFP+ cells co-stained with the proliferation marker Ki67 (Scale bar: 100 μm).

To overcome this limitation, we identified Notch3 as a more specific marker of both normal and tumor pericytes and confirmed its cell-specific expression in AOM/DSS tumors (Figure 2E-F). Subsequently, we employed the Notch3-CreERT2 strain (*42*) in combination with mTmG mice and performed lineage tracing using the same protocol described above (Figure 2G). The Notch3-CreERT2 mice specifically targeted MCAM+ pericytes in the normal colon. These labeled pericytes showed a significant increase in AOM/DSS tumors (Figure 2H-J). Immunostaining for Ki67 revealed proliferating GFP+ perivascular cells in AOM/DSS tumors in both PDGFRβ-CreERT2;mTmG and Notch3-CreERT2;mTmG mice (Figure 2K). These results demonstrate that pericytes proliferate and increase in numbers during colitis-associated carcinogenesis.

### Notch3 signaling regulates pericyte proliferation and blood vessel properties in the intestinal TME

In addition to identifying Notch3’s restricted expression in pericytes, we next investigated if it also had a functional role. We initially performed cell-to-cell interaction analysis using CellChat (*43*), which revealed that pericytes are a major target (incoming signal strength) and source (outgoing signal strength) of the Notch pathway (Figure 3A). The Notch pathway was particularly active in the interactions in-between pericytes (autocrine signaling) and between pericytes and endothelial cells (paracrine signaling) (Figure 3B). Among the different ligand-receptor pairs in the TME, the Jag1-Notch3 and Dll4-Notch3 pairs contribute significantly to this enrichment (Figure 3C). Notch3 was expressed in tumor pericytes and accepted signals from pericytes (*Jag1*) and blood endothelial cells (*Jag1*, *Dll4*), including proliferating endothelial cells (Figure 3D-F). Notably, the downstream Notch3 target Heyl was also restrictively expressed in pericytes, further supporting the cell-specific activation of the pathway (Figure 3D). These findings support the activation of Notch3 signaling and its potential regulation of pericyte function within the intestinal TME.

**Figure 3.**
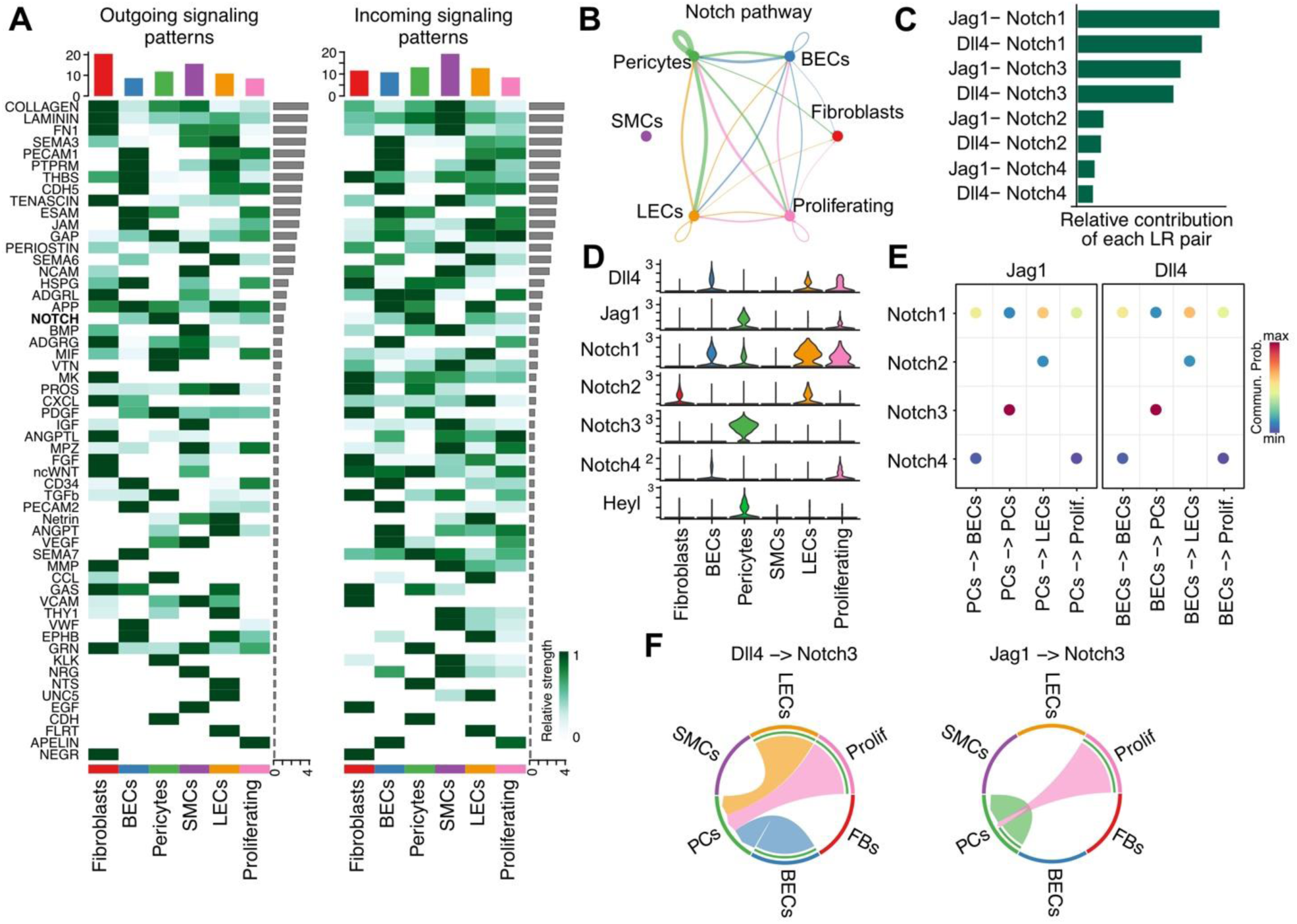
Ligand-receptor interaction analysis in the AOM/DSS tumor stroma. (A) Heatmap showing the signaling pathways that contribute to outgoing or incoming communication. The color bar represents the relative signaling strength of a signaling pathway across cell types. The bars indicate the sum of the signaling strength of each cell type or pathway. (B) The inferred NOTCH signaling network. The edge weights represent the aggregated interaction strength of all ligand-receptor pairs of the NOTCH signaling pathway. (C) The contribution of each ligand-receptor pair to the overall signaling of the inferred NOTCH pathway. (D) Violin plots showing the expression distribution of signaling genes involved in the inferred Notch3 signaling network in the AOM/DSS tumor stroma. (E) Bubble plot showing inferred interactions between Notch ligands and receptors in selected cell groups (p < 0.01). (F) Chord plots showing the inferred Dll4-Notch3 and Jag1-Notch3 signaling networks in the AOM/DSS tumor stroma. FBs, fibroblasts; PCs, pericytes. All inferred interactions are statistically significant (p.adj ≤ 0.05).

To investigate the pathophysiological role of Notch3 in intestinal tumors, we utilized a mouse strain that enables the inducible expression of the intracellular domain of Notch3 upon Cre-mediated recombination, leading to the cell-specific activation of the pathway (*44*). We crossed these mice with the PDGFRβ-CreERT2 strain and confirmed increased Notch3 expression in isolated stromal cells following tamoxifen administration (Notch3ICD or N3ICD) (Figure 4A and Supplementary Figure 4). We then applied the AOM/DSS protocol to these mice and their littermate controls, all of which received five consecutive tamoxifen injections one week before the initiation of colitis-associated carcinogenesis (Figure 2A). While mice did not display statistically significant differences in tumor multiplicity or size (Figure 4B-C), tissue staining with antibodies against Ki67 and αSMA, revealed increased proliferation of GFP+ cells and reduced expression of αSMA expression around blood vessels (Figure 4D-E). These results suggest that Notch3 plays a role in balancing proliferation and activation of pericytes. To verify the effect of Notch3 pathway activation on pericyte properties, we utilized the Rosa26-CreERT2 mouse (*45*), which we crossed with N3ICD mice. We FACS-sorted EpCAM-CD45-MCAM+ cells from these mice and incubated them with 4-hydroxytamoxifen (4-OHT) to induce Notch3 activation (Figure 4F). Cells from Rosa26-CreERT2;mTmG mice were used to confirm the efficiency of this method (Figure 4G). MTT assay measurements demonstrated increased proliferation in cells with activated Notch3 signaling, while other growth factors, including TGFβ and PDGF, did not have a pro-proliferative effect either alone or in the presence of Notch3 activation (Figure 4H). Therefore, our results show that Notch3 signaling promotes pericyte proliferation in vitro and vivo.

**Figure 4.**
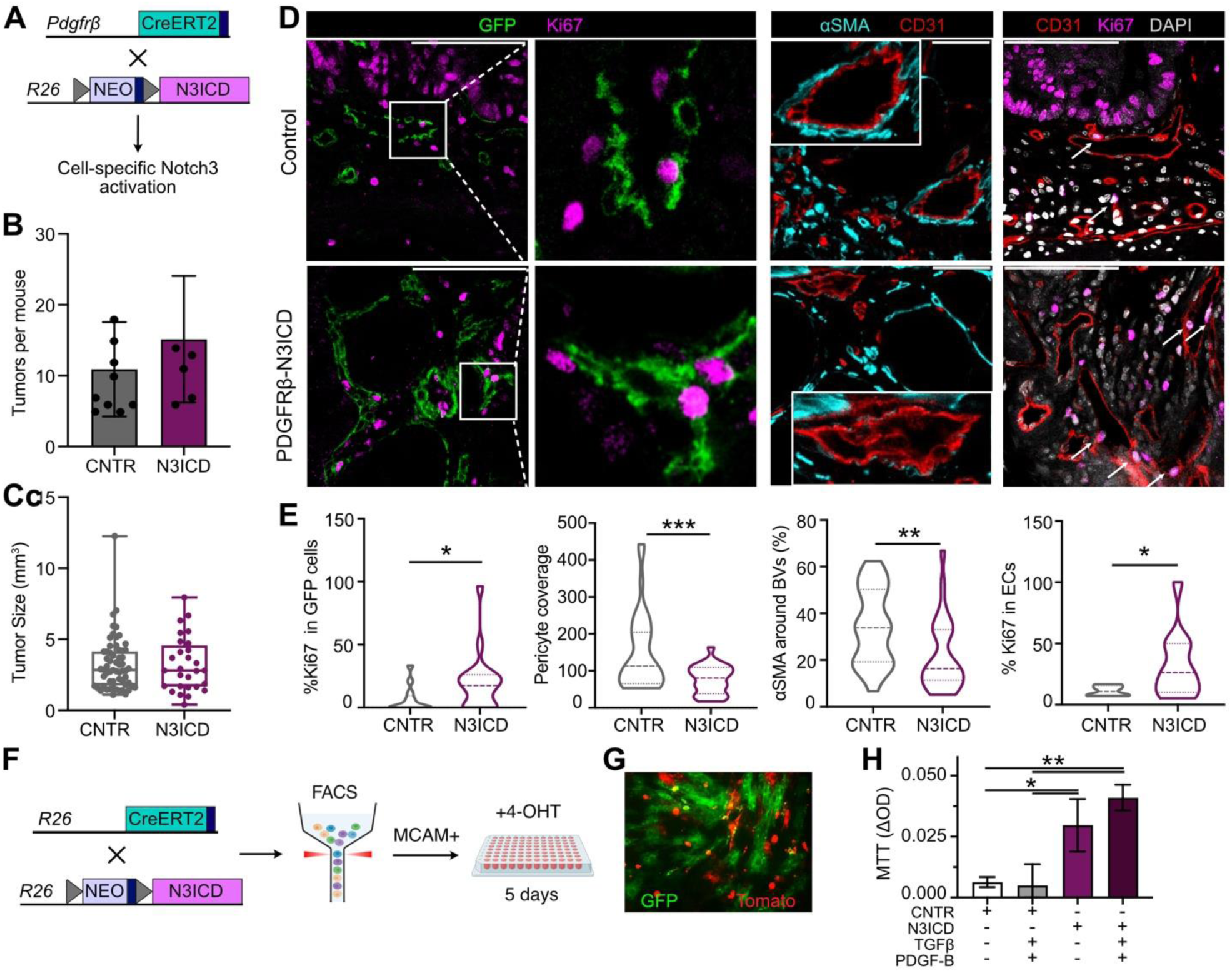
Stromal Notch3 activation regulates pericyte and endothelial cell properties. (A) Schematic representation of the in vivo strategy for cell-specific Notch3 pathway activation. (B) Number of tumors per mouse and (C) mean tumor size in Notch3-activated mice (N3ICD) and their littermate wildtype controls (CNTR) (one of two experiments, unpaired t-test). (D) Representative confocal images and (E) quantifications showing proliferative GFP (Ki67+GFP+) cells (n = 13-19, scale bar: 100 μm (up) and magnifications (down)), αSMA expression around blood vessels (BVs) (n = 26-29 BVs, scale bar: 20 μm), pericyte coverage (n = 26-40 BVs), and proliferative endothelial cells (Ki67+CD31+) cells (n = 10-20, scale bar: 100 μm, white arrows indicate ki67+ endothelial cells) in AOM/DSS tumors from N3ICD mice and littermate controls (n = 7-10 tumors from one experiment out of two). Results were analyzed using Mann-Whitney test and unpaired two-tailed t-test. (F) Schematic representation of in vitro activation of Notch3 in MCAM+ cells and assessment of cell survival (partly generated using Biorender.com). (G) GFP expression in MCAM+ cells isolated from R26CreERT2-mTmG mice following 5 days incubation with 4-hydroxitamoxifen (4-OHT). (H) Results from the MTT assay, showing the proliferation of MCAM+ cells that have active Notch3 signaling, in the presence or absence of the growth factors TGFβ and PDGF-B (n = 3 from one of two experiments). Results were analyzed using one-way ANOVA (*p < 0.05, **p < 0.01, ***p < 0.001).

Notably, we also observed increased CD31+ cell proliferation as a consequence of Notch3 activation in PDGFRβ+ cells (Figure 4D-E). However, despite the parallel increased proliferation of both cell types, pericyte coverage, expressed as the ratio of % GFP+ to CD31+ cells around blood vessels, was reduced in PDGFRβ-N3ICD tumors in comparison to controls (Figure 4D-E). These results suggest that increased pericyte proliferation downstream of Notch3 activation is associated with increased endothelial cell proliferation and reduced pericyte coverage, which is indicative of disrupted vessel integrity inside CAC tumors (*8, 9, 46*).

### Notch3 deletion results in pericyte phenotype changes hindering cancer progression

To better understand the pathophysiological significance of pericyte-specific Notch3 signaling in intestinal carcinogenesis, we also utilized Notch3 deficient mice (*47*). After applying the AOM/DSS protocol of CAC, we observed no difference in tumor multiplicity and size compared to controls, similar to our observation with Notch3 activation (Figure 5A-B). However, using similar staining to our previous experiments, we found that Notch3 deletion led to decreased proliferation of CD31+ endothelial cells (Figure 5C-D). αSMA staining around blood vessels was similar between experimental and control mice, suggesting that Notch3 activation may be sufficient but not necessary for pericyte activation in vivo (Figure 5C-D). Accordingly, pericyte coverage, was not altered (Figure 5C-D).

**Figure 5.**
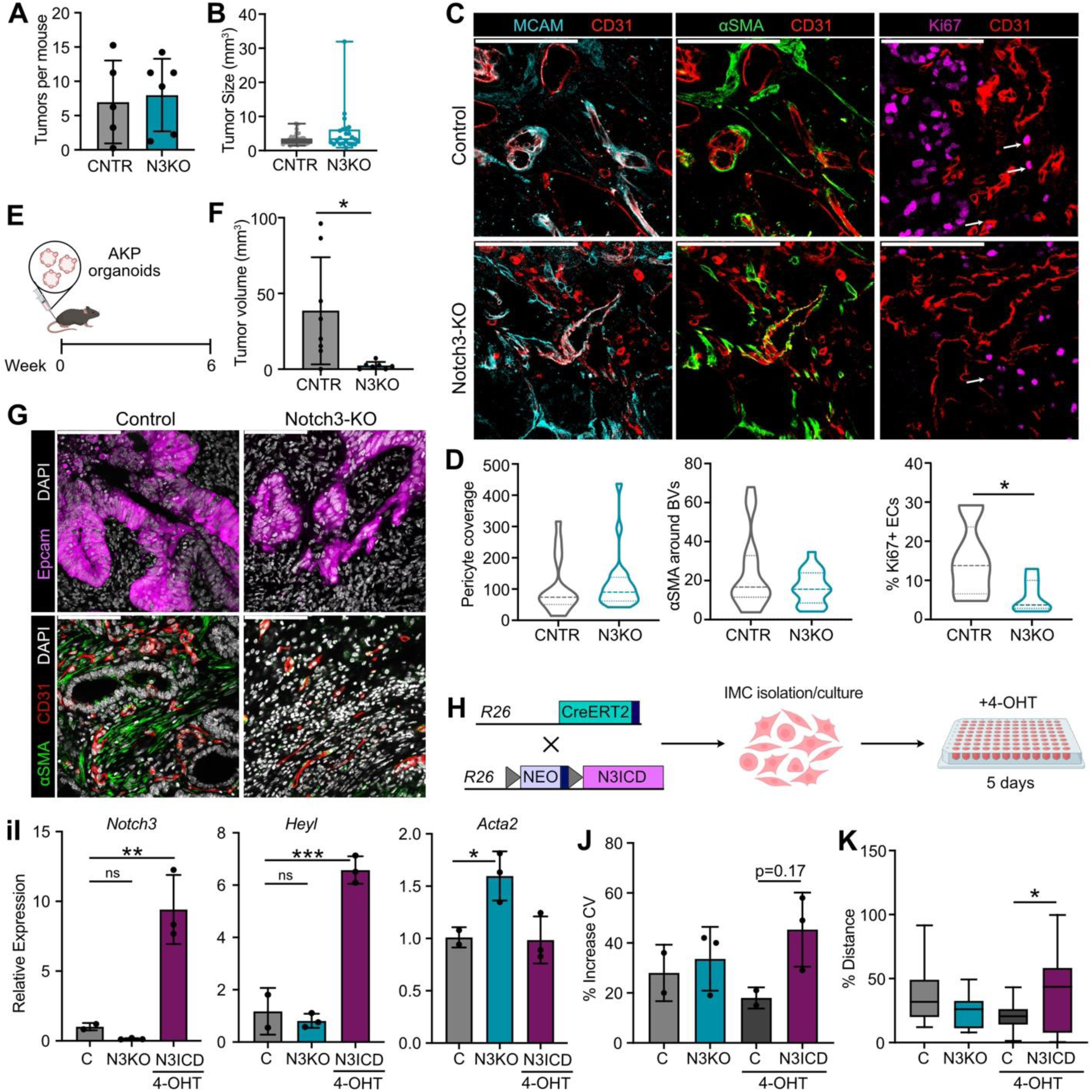
Notch3 deletion inhibits colorectal cancer progression. (A) Number of tumors per mouse and (B) mean tumor size in Notch3 knockout mice (N3KO) and their littermate wildtype controls (CNTR) (one of two experiments, unpaired t-test) (C) Representative IHC images and (D) quantifications showing αSMA expression around blood vessels (n = 18-19 BVs), proliferative endothelial cells (Ki67+CD31+) cells (n = 7-9 BVs), and pericyte coverage (MCAM-CD31 staining) (n = 17-20 BVs). Results were analyzed using unpaired 2-tailed t-test. Scale bar: 100 μm. (E) Schematic representation of the AKP mucosal injection model (partly generated using Biorender.com). (F) Size of AKP tumors in Notch3 deficient mice and littermate controls (n = 7-8 mice, Welch’s t-test). (G) Representative IHC images showing epithelial cells (EpCAM+), endothelial cells (CD31+) and αSMA expression in AKP tumors (scale bar: 100 μm). (H) Schematic representation of in vitro activation/deletion of Notch3 in fibroblasts (partly generated using Biorender.com). (I) Gene expression analysis of *Notch3*, *Heyl*, and *Acta2* in isolated stromal cells with Notch3 deletion (N3KO) or activation (N3ICD) (n = 2-3 biological replicates). (J) Quantification of the differences in crystal violet staining (CV) 24 hours after plating of the stromal cells, and (K) Quantification of the distance in wound healing assay (n = 2-3 biological replicates, results were analyzed using one-way ANOVA). Notch3 activation was induced in isolated Rosa-CreERT2;N3ICD intestinal stromal cells after incubation with 4-hydroxytamoxifen (4-OHT) for 5 days.

Since the AOM/DSS model primarily results in adenomas and the integrity of blood vessels in tumors plays a crucial role in metastasis, we also investigated whether Notch3 deletion could influence later stages of tumor progression. For this purpose, we utilized an orthotopic transplantation model in which AKP organoids – bearing deletion of *Apc*, *Tp53*, along with an activating mutation in *Kras* – were injected into the mouse rectum (*48*) (Figure 5E). Six weeks after transplantation, all wildtype control mice had developed tumors ranging in size from 12 to 96 mm^3^. However, tumors were either absent or significantly smaller in the Notch3-KO mice (Figure 5F). Stainings revealed fewer and smaller blood vessels in Notch3-KO tumors compared to control tumors, albeit the small number and size of tumors did not allow for image analysis and quantifications (Figure 5G). These findings indicate that Notch3 signaling plays a crucial role in advanced stages of colorectal cancer progression.

To further investigate the opposite functions of Notch3 deletion and activation, we performed in vitro experiments, similar to those in Figure 4f-h (Figure 5H). To overcome the challenge of low MCAM+ cell yield, we used total stromal cells isolated from Notch3 knockout mice (N3KO), Rosa-CreERT2;N3ICD (N3ICD) mice, and controls (C, WT or N3ICDf/f). We confirmed increased expression of the intracellular domain of Notch3 in N3ICD cells, and the concomitant increase in the Notch3-target gene *Heyl* (Figure 5I). Notch3 expression was reduced in Notch3 knockout cells, although the difference was not statistically significantly, possibly due to the relatively low number of Notch3-expressing pericytes in the control cultures (Figure 5I). However, *Acta2* was increased in the absence of *Notch3*, suggesting a negative association, which aligns with our previous in vivo observations (Figure 4D-E). Notch3 activation also led to an increase – albeit not statistically significant - in cell proliferation, as assessed by crystal violet staining, and reduced wound closure ability (Figure 5J-K). These results suggest that Notch3 activation may promote pericyte proliferation while limiting their contractile activation.

### Cancer-associated pericyte phenotypic states are regulated by Notch3 signaling

Considering the in vivo effects of Notch3 activation/deletion on pericyte properties, we investigated in detail the heterogeneity of cancer-associated pericytes in our dataset. We performed subclustering of 1008 pericytes (mainly originating from the AOM/DSS sample – 1000/1008 cells) and analyzed the transcriptomic profiles of 4 subsets, which differed in the expression level of genes associated with ECM production and remodeling, myogenic features, inflammatory activation, and endothelial traits. The two largest subsets exhibited classic markers of pericytes (e.g. *Notch3, Pdgfrb, and Ndufa4l2*), but different levels of *Rgs5* and ECM-related molecules (*Col3a1*, *Vtn, Mmp11*), and myogenic-related genes (*Acta2*, *Myh11*, *Tagln*). As such, we designated them ecmPCs and myPCs, respectively. A smaller cluster expressed genes related to inflammatory activation (*Mmp10*, *Saa3*, *Cxcl1*, *Il6*), and we labeled it iPCs. Finally, there was a distinct cluster expressing endothelial cell markers, such as *Pecam1*, *Plvap, Cdh5,* and *Kdr*, which indicates endothelial to mesenchymal transition, as previously described (endoPCs) (Figure 6A-B and Supplementary Data 1) (*31*). The presence of these subsets was also verified by FACS analysis in both normal and tumor samples, although iPCs were not detectable in the normal colon (Figure 6C-D). The relative abundance of the other three subsets was not statistically significantly different between the normal colon and AOM/DSS induced adenomas (Figure 6C-D).

**Figure 6.**
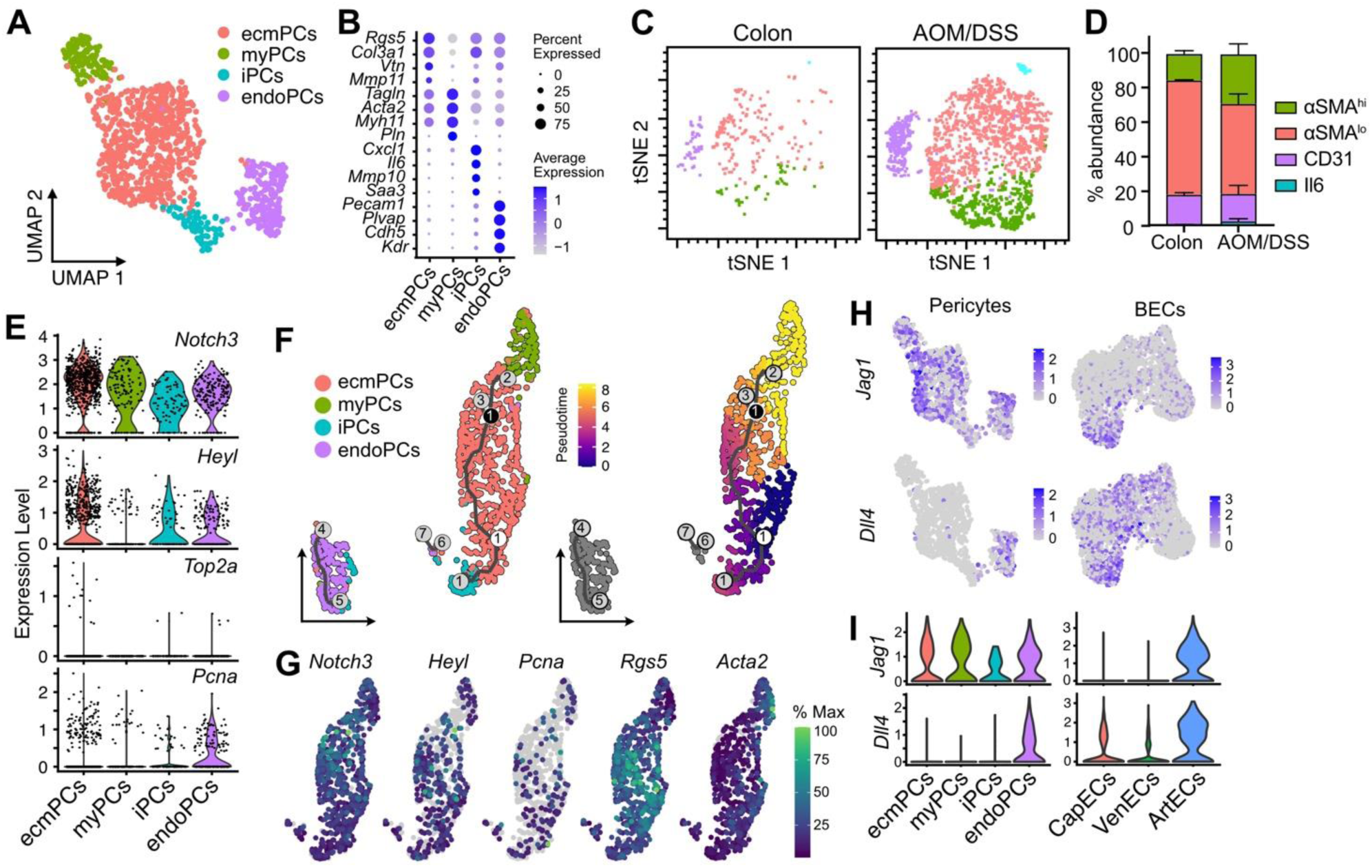
Pericyte heterogeneity in colitis-associated cancer. (A) UMAP analysis of AOM/DSS tumor pericytes, colored by pericyte cluster. (B) Dot plot showing marker genes for each pericyte cluster (p.adj ≤ 0.05). (C) FACS tSNE plots and (D) quantification of MCAM+αSMA+ pericyte subpopulations in the normal colon and CAC tumors. (E) Violin plots showing expression of genes associated with Notch3 signaling and markers of proliferation in each pericyte cluster. (F) Trajectory analysis showing potential transition between pericyte clusters and in pseudotime. (G) Expression of selected genes in the monocle3 UMAPs. (H) Feature and (I) violin plots showing expression of Notch3 ligands *Jag1* and *Dll4* in pericyte and blood endothelial subsets.

Examining the expression of *Notch3*, the Notch target gene *Heyl*, and proliferation markers *Top2a*, and *Pcna*, we found that ecmPCs were enriched in *Heyl*, which was absent in myPCs. Proliferation markers were also increased in ecmPCs, in agreement with our vivo and in vitro data (Figure 6E). Trajectory analysis using monocle 3 (*49*) showed a possible transition from ecmPCs to myPCs, which correlated with decreased *Heyl* expression (Figure 6F-G). These results suggest that pericytes can be found in distinct phenotypic states that are regulated by Notch3 signaling, instructed by direct cell-to-cell contact with other pericytes or Notch3-ligand expressing endothelial cells, and, therefore, influenced by cell topology. Among endothelial subsets, capillary endothelial cells are the main cellular source of both Jag1 and Dll4 ligands in the TME (Figure 6H-I).

We next validated these results in our combined human colorectal cancer dataset through subclustering of pericytes. We observed the presence of myPCs, and two clusters of ecmPCs, one enriched in tumors (T) and the other in the normal tissue (N). Interestingly, an intermediate cluster between myPCs and tumor ecmPCs was also evident (Figure 7A-D and Supplementary Data 1). We also detected a small cluster of proliferating PCs (prolifPCs) in tumors (Figure 7A-D). Inflammatory pericytes, although present, did not form a distinct subset (Figure 7E). It should be noted that endoPCs were clustered together with blood endothelial cells and as such they were not included in the subclustering (Figure 7F). Trajectory analysis using monocle 3 showed the potential of normal ecmPCs as a source of both tumor ecmPCs and myPCs (Figure 7G). Interestingly, *HEYL* was preferentially expressed in tumor ecmPCs, intPCs, and prolifPCs, consistent with our mouse data (Figure 7C, H). In conclusion, our analysis supports a model where Notch3 activation in response to microenvironmental cues, drives pericyte proliferation, a shift to synthetic ecmPC state, and subsequent endothelial proliferation. Conversely, Notch3 deletion, associated with contractile myPCs, leads to reduced endothelial cell proliferation, and potential vascular normalization, hindering cancer progression (Figure 7I).

**Figure 7.**
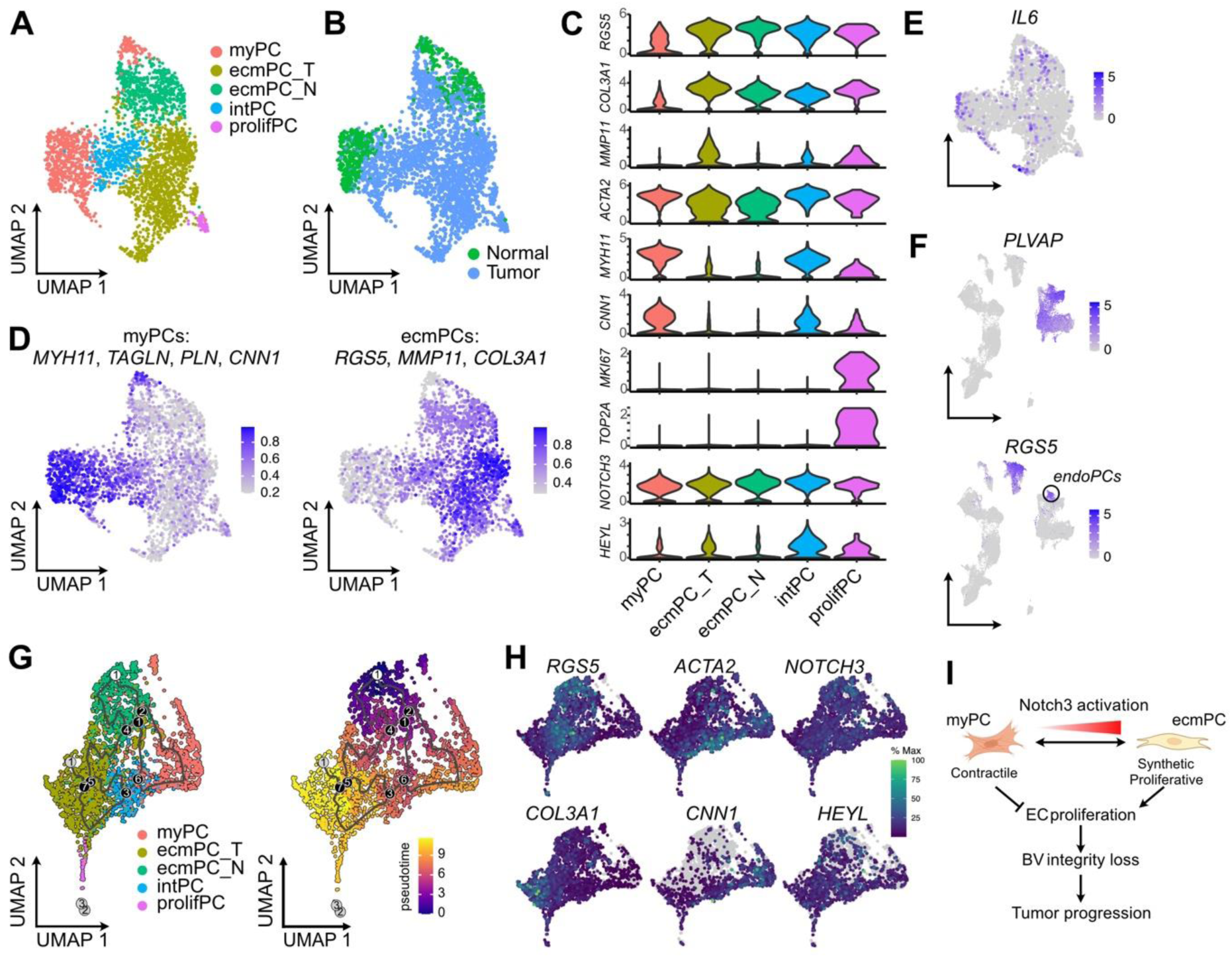
Pericyte heterogeneity in human colorectal cancer. UMAP analysis of normal and tumor pericyte in colorectal cancer patients, colored by (A) pericyte cluster, (B) tissue of origin. (C) Violin plots showing marker gene expression for pericyte clusters, as well as *NOTCH3* and *HEYL*. (D) Feature plots showing expression of a myogenic and an ECM-related signature. Feature plots showing expression of (E) IL6 in human pericytes and (F) *PLVAP* and *RGS5* in the human colorectal cancer stroma. The circle denotes cells that express both endothelial (*PLVAP*) and pericyte (*RGS5*) markers corresponding to endoPCs. (G) Trajectory analysis showing transition between pericyte clusters and in pseudotime. (H) Expression of selected genes in the monocle3 UMAPs. (I) Schematical representation of proposed Notch3 function in intestinal cancer. (J) Schematic representation of Notch3’s proposed function in the regulation of pericyte plasticity and function in CRC (partly generated using Biorender).

## Discussion

Our study provides several novel insights into the role of pericytes in CRC. These cells, often referred to as cancer-associated pericytes (CAPs) or vascular CAFs (vCAFs) have received considerably less attention in comparison to other TME components, despite their crucial functions (*50*). First, we showed that pericytes are enriched in the AOM/DSS tumor microenvironment and actively proliferate within tumors, contributing to the altered stromal composition. This observation was confirmed in published human CRC datasets, highlighting the consistency of our results across species and the relevance to human CRC. Previous reports have suggested significant contributions from bone marrow-derived cells to the pericyte pool in pancreatic and brain tumors (*11, 12*). Given that bone marrow cells do not contribute to the AOM/DSS stroma (*34*), normal resident pericytes seem to be the primary source of tumor pericytes. In addition, despite previous reports suggesting the presence of resident pericyte progenitors (*51, 52*), we did not identify a specialized pericyte subset with a progenitor expression profile. However, we did detect pericytes with endothelial characteristics, indicative of endothelial-to-mesenchymal transition (EndoMT) (*13, 31*). Nevertheless, their equal contribution to both the normal tissue and tumor microenvironment along with their distinct signature points towards their proliferation rather than complete transdifferentiation. The different cellular sources reported for tumor pericytes underline the context-dependent nature of tumor pericyte recruitment across different cancer types and experimental models. The identification of resident pericytes as a primary source of tumor pericytes in CRC suggests that interventions targeting pericyte function rather than recruitment might be more effective in this context.

Second, we identify Notch3 as a key regulator of pericyte function in CRC, influencing their proliferation, activation, and interaction with endothelial cells. This pro-proliferative effect of Notch3 signaling in pericytes is consistent with its role in vascular smooth muscle cells, where Notch3 activation has been shown to enhance proliferation and inhibit apoptosis (*20, 53*). Accordingly, in vitro activation of Notch3 signaling inhibited vSMC migration and led to downregulation of contractile gene expression (*53, 54*). The dual role of Notch3 in promoting pericyte proliferation while limiting their contractile activation has important implications for tumor angiogenesis and metastasis. Reduced pericyte coverage and activation around tumor vessels can lead to increased vascular permeability, impaired blood flow, and enhanced tumor cell intravasation, facilitating metastasis (*15*). Our finding that Notch3 deletion impairs tumor growth in an orthotopic model of advanced CRC suggests that inhibiting Notch3 signaling may be beneficial in treating advanced stages of the disease by normalizing the tumor vasculature and limiting metastatic spread.

Third, we reveal the heterogeneity of CRC-associated pericytes and show that Notch3 signaling may regulate transitions between different pericyte phenotypes or states. The identification of distinct pericyte subpopulations with different functional properties in both mouse and human CRC tumors highlights the complexity of the tumor microenvironment. These findings are consistent with emerging evidence for pericyte heterogeneity in both normal tissues and pathological conditions, including cancer (*31*). Notably, the identified pericyte subpopulations exhibited differential expression of Notch pathway components and target genes, suggesting that Notch3 signaling may influence pericyte heterogeneity and plasticity in the tumor microenvironment. We specifically identified a pericyte subpopulation with high Notch3 activity and low expression of contractile proteins, and a contractile subpopulation with low Notch3 activity and high expression of αSMA and other contractile markers. This observation supports our functional studies demonstrating Notch3-mediated regulation of pericyte proliferation and contractility and suggests that these phenotypic states represent distinct positions along a continuum of pericyte differentiation. This result also fits into the concept of phenotype switching, a process characterized by pericyte transition between different states, such as from a quiescent, contractile phenotype that stabilizes blood vessels to a more plastic, proliferative, or angiogenic state associated with tissue remodeling, disease progression, or metastasis (*8–10*). Indeed, chemical inhibition of Notch using a γ-secretase inhibitor was previously shown to drive the generation of a more contractile and quiescent pericyte phenotype (*9*). The regulation of these phenotypic states by Notch3 signaling provides a potential mechanism for therapeutic intervention, targeting specific pericyte subpopulations to modulate the tumor microenvironment.

Finally, we demonstrate that targeting Notch3 signaling in pericytes can impair tumor growth in advanced CRC models, suggesting a potential therapeutic approach. Traditional anti-angiogenic therapies focusing on VEGF/VEGFR signaling have shown limited efficacy in CRC (*15, 55*), highlighting the need for alternative approaches. Targeting the Notch3 pathway represents a novel strategy that could complement existing anti-angiogenic therapies by promoting vessel normalization. Based on our results, inhibition of Notch3 signaling in pericytes would be expected to reduce pericyte proliferation and enhance their contractile phenotype, potentially leading to improved vessel stability and reduced leakiness. This normalization effect could enhance drug delivery to tumors and reduce hypoxia-driven tumor aggressiveness (*55*). A similar effect was recently shown with the use of MEK, PI3K, or general Notch inhibition (*9*). Furthermore, normalized vasculature may facilitate immune cell infiltration, potentially enhancing the efficacy of immunotherapies, which have shown limited success in CRC (*3, 9, 10*). Conversely, targeting a specific Notch ligand in contrast to general Notch inhibition may also lead to reduced potential side-effects.

In conclusion, our study shows that pericytes are enriched in the CRC microenvironment and that Notch3 signaling plays an important role in regulating their proliferation, activation, and interaction with endothelial cells. The heterogeneity of tumor-associated pericytes and their regulation by Notch3 signaling provide novel insights into the complexity and plasticity of the tumor microenvironment. Targeting Notch3 signaling in pericytes to modulate pericyte phenotype may represent a promising therapeutic approach for advanced colorectal cancer and other malignancies.

## Materials and Methods

### Animals

PDGFRβ-CreERT2 (*39*), Notch3-/- (*47*), and Rosa26-CreERT2 (*45*) were purchased by Jackson Laboratories. Notch3-CreERT2 (*42*), Rosa26-N3IC (*44*) mice were a kind gift from S. Artavanis-Tsakonas (Harvard Medical School, Harvard University, Boston), S. Fre (Institut Curie, CNRS, Paris), and I. Aifantis (NYU School of Medicine, New York). All mice were maintained on the C57Bl/6 background. Eight to sixteen-week-old male and female mice were used for all experiments. They were housed in the animal house facility of the Biomedical Sciences Research Center Alexander Fleming under specific pathogen-free conditions with controlled temperature (22 ± 2°C), humidity (55 ± 10%), and a 12-hour light/dark cycle. All animal experiments were approved by the Institutional Animal Care and Use Committee of BSRC Fleming (protocol numbers: 1439373, 1439369, 68130) and conducted in accordance with European and national guidelines for the care and use of laboratory animals.

### Tamoxifen injections

For inducible Cre-mediated recombination, mice were injected intraperitoneally (i.p.) with 1mg of tamoxifen (Sigma) per mouse daily for five consecutive days. Tamoxifen was diluted in corn oil at 10mg/ml.

### AOM/DSS-induced colitis-associated cancer (CAC) model

Colitis-associated cancer was induced using the azoxymethane (AOM)/dextran sodium sulfate (DSS) model as previously described (*29*). Briefly, mice were injected intraperitoneally with a single dose of AOM (10 mg/kg body weight; Sigma-Aldrich) on day 0. One week after AOM injection (day 7), mice were administered 2.5% DSS (molecular weight 36,000-50,000 Da; MP Biomedicals) in the drinking water for 5 consecutive days, followed by 16 days of regular drinking water. This DSS treatment cycle was repeated twice more for a total of three cycles. Mice were monitored daily for body weight, stool consistency, and rectal bleeding. Animals were sacrificed on day 60 after AOM injection by CO₂ inhalation followed by cervical dislocation. The entire colon was removed, flushed with ice-cold PBS, and opened longitudinally. The number, size, and location of visible tumors were recorded. The colon was then fixed in 4% PFA for 4-6 hours and processed for agarose embedding and vibratome sectioning.

### AKP mucosal injection model

The AKP mucosal injection model was performed as previously described (*19*). Briefly, mouse AKP organoids (a kind gift from O. Yilmaz, Koch Institute, MIT) were embedded in Matrigel (Corning), and cultured in Advanced DMEM/F12 (supplemented with 2mM GlutaMAX (ThermoFisher Scientific), 10mM HEPES (ThermoFisher Scientific), 1X B-27 supplement (ThermoFisher Scientific), 1X N-2 supplement (ThermoFisher Scientific), 1mM N-acetyl-L-cysteine (Sigma), 10μM Y-27632 (Cell Signaling Technology), 10μM Nutlin-3a (Cayman), 100ng/ml Noggin (Peprotech), 100U/ml Penicillin, 100mg/ml Streptomycin ((ThermoFisher Scientific), and 50μg/ml Gentamycin (Applichem)). Prior to injections, AKP organoids were dissociated using TrypLE Express (ThermoFisher Scientific) for 1min at 37°C and resuspended in Advanced DMEM, containing 2mM L-Glutamine (Biosera), 1X B-27 supplement, 1X N-2 supplement, and 10 μM Y-27632, with 10% Matrigel (Corning). Organoids were subsequently injected directly into the colon mucosa of mice. Each injection contained 500 organoids in 30 μl of the resuspension medium. Tumors were collected 6 weeks after injection and their size was measured using a digital caliper.

### Immunohistochemistry

Tissues embedded in 3% low melting agarose (Lonza) were sectioned at 80 μm thickness using a vibratome (Leica Biosystems). After blocking using 1% BSA in PBS containing 0.05%Tween 20 (Sigma), sections were incubated with primary antibodies in blocking solution overnight at 4°C. After washing, sections were incubated with the corresponding secondary antibodies (1:500, Invitrogen) for 1 hour at room temperature. All antibodies are listed in Table 1. Nuclei were counterstained with DAPI. Images were acquired using a confocal microscope (Leica TCS SP8 White Light Laser confocal system). Quantifications were performed using ImageJ (NIH, Bethesda, MD, USA) according to the following parameters:

– Pericyte coverage was quantified based on the hypothesis that pericytes (PDGFRβ-GFP+ in N3IC samples and MCAM+ in N3KO samples) closely associate with and ensheathe endothelial cells, contributing to vessel coverage. Quantification of pericytes and endothelial cells was performed separately for the same images using the Mexican Hat filter (scale parameter: σ = 2 px). The filter was applied to enhance circular structures, emphasizing signal detection while suppressing high-frequency noise. The area fraction (%) of the pericyte-specific signal and the CD31+ endothelial signal was measured independently for each image. Pericyte coverage was then expressed as the ratio of pericyte-positive area (%) to CD31-positive area (%) per field of view, providing a normalized measure of pericyte investment relative to the endothelial surface.
– For Quantification of αSMA+ signal around blood vessels, vessel segments were selected based on the presence of CD31+ endothelial staining without DAPI+ nuclei within the lumen, ensuring analysis of vessel structures devoid of intraluminal cells. Using the freehand selection tool, regions of interest (ROIs) were manually drawn to encompass each vessel and its surrounding αSMA+ signal. The ROIs were converted to 8-bit grayscale and applied a Mexican Hat filter (σ = 2 px) to enhance tubular structures. Following this preprocessing, the area directly covered by αSMA staining within the initial ROI was delineated using the freehand selection tool, and the mean gray value of the αSMA signal was measured. This approach allowed for the assessment of αSMA coverage intensity around individual vessel segments.
– Proliferating endothelial (CD31+Ki67+) and perivascular (PDGFRβ+Ki67+) cells were quantified as follows: for each field of view, the total number of DAPI+ nuclei within CD31+ endothelial cells or PDGFRβ+ perivascular cells was manually counted. For vessel segment analysis, selection was based on the presence of CD31+ endothelial staining without DAPI+ nuclei within the vessel lumen, ensuring analysis of vessel structures devoid of intraluminal cells. Subsequently, Ki67+ nuclei co-localizing with DAPI+ nuclei within the CD31+ or PDGFRβ+ cell populations were counted. The percentage of proliferating cells was calculated by dividing the number of Ki67+CD31+ or Ki67+PDGFRβ+ cells by the total number of CD31+DAPI+ or PDGFRβ+DAPI+ cells, respectively, per field of view.

**Table 1.**
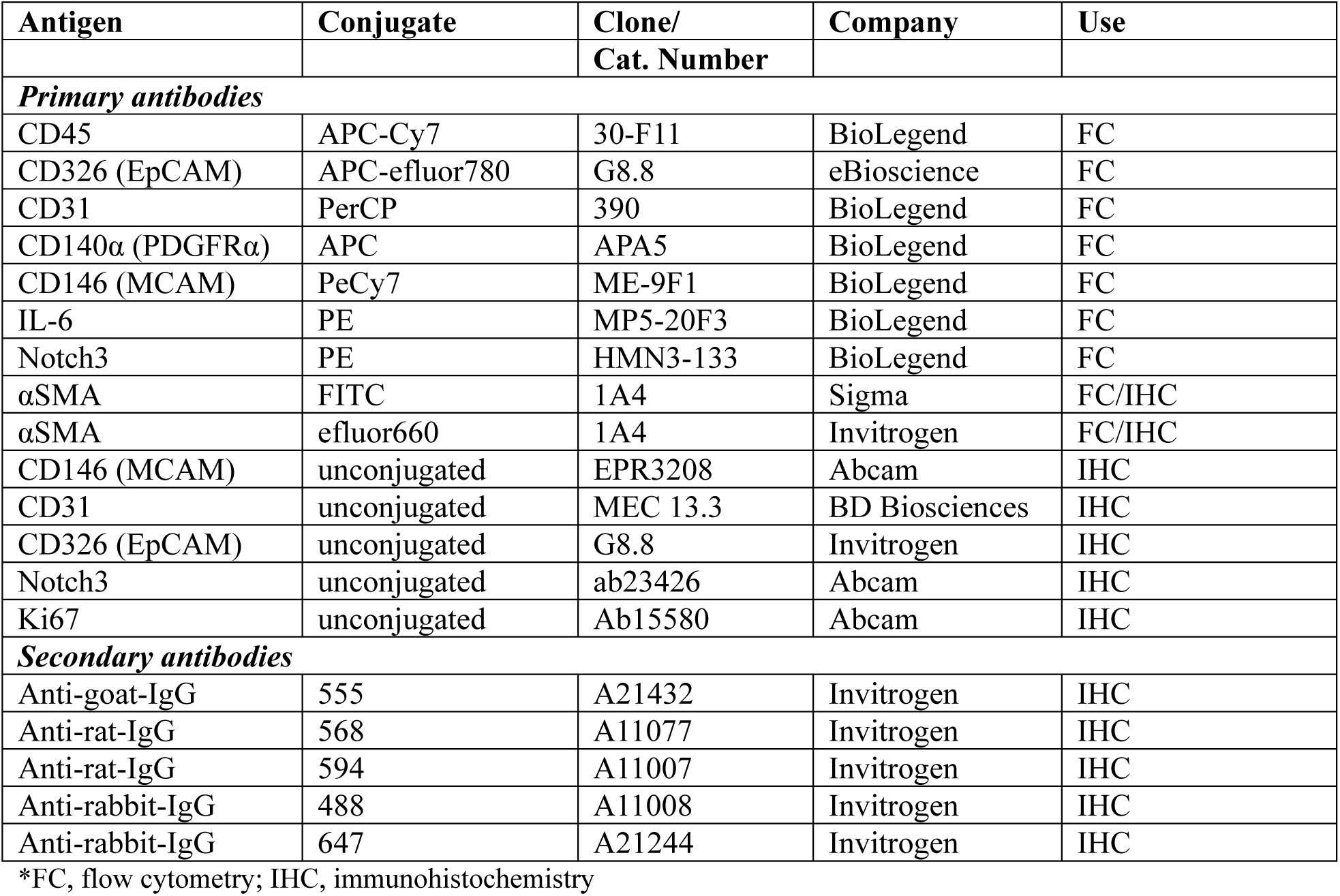
List of antibodies used in immunohistochemistry and flow cytometry.

| Antigen | Conjugate | Clone/ | Company | Use |
| --- | --- | --- | --- | --- |

|  |  | Cat. Number |  |  |
| --- | --- | --- | --- | --- |
| **Primary antibodies** |  |  |  |  |
| CD45 | APC-Cy7 | 30-F11 | BioLegend | FC |
| CD326 (EpCAM) | APC-eFluor780 | G8.8 | eBioscience | FC |
| CD31 | PerCP | 390 | BioLegend | FC |
| CD140α (PDGFRα) | APC | APA5 | BioLegend | FC |
| CD146 (MCAM) | PeCy7 | ME-9F1 | BioLegend | FC |
| IL-6 | PE | MP5-20F3 | BioLegend | FC |
| Notch3 | PE | HMN3-133 | BioLegend | FC |
| αSMA | FITC | 1A4 | Sigma | FC/IHC |
| αSMA | eFluor660 | 1A4 | Invitrogen | FC/IHC |
| CD146 (MCAM) | unconjugated | EPR3208 | Abcam | IHC |
| CD31 | unconjugated | MEC 13.3 | BD Biosciences | IHC |
| CD326 (EpCAM) | unconjugated | G8.8 | Invitrogen | IHC |
| Notch3 | unconjugated | ab23426 | Abcam | IHC |
| Ki67 | unconjugated | Ab15580 | Abcam | IHC |
| **Secondary antibodies** |  |  |  |  |
| Anti-goat-IgG | 555 | A21432 | Invitrogen | IHC |
| Anti-rat-IgG | 568 | A11077 | Invitrogen | IHC |
| Anti-rat-IgG | 594 | A11007 | Invitrogen | IHC |
| Anti-rabbit-IgG | 488 | A11008 | Invitrogen | IHC |
| Anti-rabbit-IgG | 647 | A21244 | Invitrogen | IHC |

\*FC, flow cytometry; IHC, immunohistochemistry

### Flow cytometry analysis and cell isolation

Flow cytometry analysis and isolation was performed as previously described (*56, 57*). Briefly, the mouse colon or AOM/DSS tumors were dissected and washed with HBSS (Gibco) supplemented with antibiotic antimycotic solution (Gibco). Epithelial cells were removed by incubating the tissue with 5 mM EDTA and 1 mM DTT in HBSS for 25 minutes at 37°C. The colonic tissues were digested in 300 U/mL Collagenase XI (Sigma-Aldrich) and 1 mg/mL Dispase II (Roche) in DMEM for 60 min at 37°C. Tumors were digested in 1000 U/mL Collagenase IV (Sigma-Aldrich, St. Louis, MO, USA), 1 mg/mL Dispase II (Roche, Basel, Switzerland), and 100 U/mL Dnase I (Sigma-Aldrich, St. Louis, MO, USA) in DMEM (BioSera, France) in three serial 20-min digestions. Cell suspensions were centrifuged and washed in FACS buffer (5% FBS (Biowest) in PBS). For stromal cell sorting, 1–2 million cells/100 μL were incubated with anti-CD45 and anti-CD326 antibodies for the exclusion of immune and remaining epithelial cells, respectively, and 1 μg/mL Propidium Iodide (PI) (Sigma) for dead cell exclusion. Cell sorting was performed using the FACSAria III cell sorter (BD) and the FACSDiva (BD) software. For pericyte sorting, cell suspensions were additional stained with an anti-CD146 antibody for positive selection and the Zombie-NIR Fixable Viability Kit (Biolegend) for selection of live cells. For stromal cell analysis, single-cell suspensions were stained with fluorochrome-conjugated antibodies against PDGFRα, MCAM/CD146, and CD31 for 30 minutes at 4°C in the dark. For intracellular stainings, cells were fixed and permeabilized using the Fixation and Permeabilization Buffer set (eBioscience), and subsequently stained with antibodies against αSMA and/or IL-6 (Table 1). Dead cell exclusion was performed using Zombie-NIR Fixable Viability Kit (Biolegend). Data were acquired on a BD Canto flow cytometer (BD Biosciences) and analyzed using the FlowJo software (version 10.9.0, FlowJo, LLC).

### Intestinal mesenchymal cell isolation

Mesenchymal cells from the mouse colon was performed as previously described (*58*). Briefly, the colon was removed and digested as described above. The cell pellet was re-suspended in culture medium, consisting of DMEM (Biochrom), 10% FBS (Biochrom), 100 U/mL penicillin/100 mg/mL streptomycin (Gibco), 2 mM L-Glutamine (Gibco), 1 μg/ml amphotericin B (Sigma) and 1% non-essential amino acids (Gibco) and plated in cell culture flasks. The medium was changed after 24h.

### RNA isolation and quantitative PCR

RNA was isolated using the RNeasy micro kit (Qiagen), cDNA was generated using the MMLV reverse transcriptase and oligo-dT primers (Promega), and quantification of gene expression was performed by qRT-PCR using the SYBR Green PCR Master Mix (Invitrogen), according to the manufacturers’ instructions. For qRT-PCR, forward and reverse primers were added at a concentration of 0.2 pmol/ml in a final volume of 20 μl for each of the following genes (Table 2). Analysis was performed on a CFX96 Touch™ Real-Time PCR Detection System (Bio-Rad).

**Table 2.**
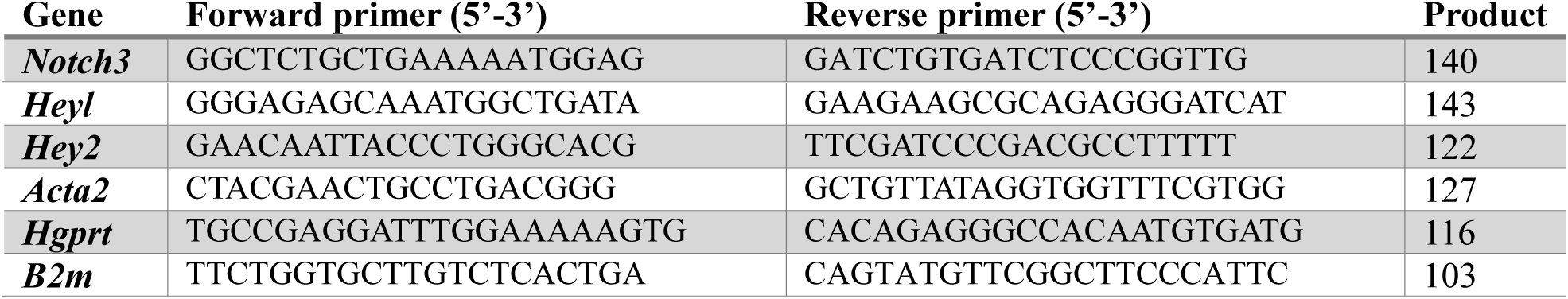
List of primers for qRT-PCR.

### In vitro wound healing assay

Cells were plated in 24-well plates and allowed to reach confluency. A 200-μl pipette tip was used to create a straight vertical “wound” in each well in the monolayer. After washing, the cultures were monitored for 24 h and the width of the wound was measured using ImageJ.

### Cell Viability Assays

Cell viability was assessed using crystal violet or the 3-(4,5-dimethylthiazol-2-yl)-2,5- diphenyltetrazolium bromide (MTT) assay. Mesenchymal cells or FACS-sorted intestinal CD146+ cells were seeded in 96-well plates at a density of 8 × 10³ cells per well in complete medium and allowed to adhere overnight. They were subsequently treated with 4- hydroxytamoxifen (4-OHT) (2μM) for 96 hours in the presence or absence of growth factors TGFβ1 (10μg/ml, Biolegend) and PDGF-B (10μg/ml, Gibco). For crystal violet staining, methanol was used for fixation and 0.1% crystal violet solution in PBS was added to the wells for 30 min. After washing, cells were left to air dry, followed by solubilization with acetic acid, and measurement of the signal at 590nm. For the MTT assay, 100 μL of MTT solution (0,5 mg/mL in complete DMEM; Sigma-Aldrich) was added to each well and incubated for 4 hours at 37°C. The medium was then removed, and 150 μL of DMSO was added to dissolve the formazan crystals. Absorbance was measured at 570 nm with a reference wavelength of 630 nm using a microplate reader (BioTek Instruments). Cell viability was calculated as a percentage relative to untreated control cells. Each experiment was performed in triplicate and repeated at least three times.

### Single-Cell Transcriptomic Analysis

#### Sample preparation

Single-cell RNA sequencing (scRNA-seq) was performed to characterize the transcriptional heterogeneity of the AOM/DSS tumor microenvironment. Fresh tumor and normal colon tissues were enzymatically dissociated as described above to obtain single-cell suspensions. Live stromal cells were isolated by FACS sorting using anti-CD45 and anti-CD326 antibodies for the exclusion of epithelial and immune cells, respectively, and propidium iodide for dead cell exclusion (Table I). Cell concentration and viability were assessed by trypan blue staining.

#### Library preparation and sequencing

For each sample, 16,500 cells were loaded onto a 10X Genomics Chromium Controller using the Chromium Next GEM Single Cell 3’ Kit v3.1 according to the manufacturer’s protocol. Library preparation was performed following the 10X Genomics workflow, with quality control performed using an Agilent Bioanalyzer High Sensitivity DNA chip. Libraries were sequenced on DNBSEQ-G400 sequencer (PE100) (BGI Genomics) with a targeted depth of approximately 200 million reads per sample.

#### Initial processing

FASTQ files were processed using the 10X Genomics CellRanger pipeline (version 7.1.0) utilizing the functionality ‘cellranger count’ with default parameters. This initial processing included alignment, Unique Molecular Identifiers (UMI) counting, filtering of empty droplets and generation of sample-specific count matrices. The number of read pairs and estimated cell numbers reached 237,166,873 bp and 6,990 cells (mean read-pairs per cell 33,929, median genes per cell 2,447) for the normal colon and 240,938,543 bp and 10,598 cells (mean read-pairs per cell 22,734, median genes per cell 1,978) for the adenocarcinomas (AOM-DSS) respectively. Read pairs were aligned to the mouse reference genome (mm10, 10X-reference genomes, 2020 version).

#### Quality control

Computational analysis was performed in R (version 4.4.2 (2024-10-31) “Pile of Leaves using the Seurat (v4) R package. Quality control was performed separately for each sample using thresholds that were manually decided based on the distribution of the quality control metrics. The metrics used were: 1) the number of unique genes in each cell (control: 700/6000, AOM/DSS: 500/7000), 2) the total number of UMIs in a cell (control: 1500/26000, AOM/DSS: 800/35000), 3) the percentage of mitochondrial gene expression (control: ≤ 10%, AOM/DSS: ≤ 15%), 4) the percentage of ribosomal gene expression (control: ≤ 3%/40%, AOM/DSS: ≤ 2%/40%), 5) the ratio between the number of expressed genes and the number of UMIs per cell (≥ 0.80), and 6) the percentage of reads mapped to the highest expressed gene for each cell (≤ 20%). Cells expressing less than 200 features were excluded.

#### Normalization, clustering, and annotation

Data normalization, identification of variable genes, gene expression scaling, dimensionality reduction, and cell clustering were performed following standard Seurat workflows for each sample. To normalize for each cell’s library size, we used the NormalizeData() function with default parameters. Cell clusters were manually annotated and merged if needed based on top cluster-specific markers and information from the literature. To compute marker genes for each cluster we used the function FindAllMarkers() excluding the features that exhibit an absolute value of average log fold change less than 0.25 between the groups of comparison and are expressed in less than 25% of the cells of each group. Doublets, as well as clusters without distinct marker genes or/and with markers corresponding to mixed cell populations accompanied also by poor quality metrics were excluded from downstream analysis. The two samples were subsequently merged using the ‘merge’ function of the Seurat package. Analysis was repeated, treating the two samples in a single object. To diminish the batch effect, we utilized the command RunHarmony from the Harmony (v1.2.3) package (theta = 2, sigma = 0.1) and manually annotated cell types, as described above. The final numbers of cells per sample were 6018 and 6674 for the control and AOM/DSS sample, respectively. Sub-clustering was performed using a similar procedure, after generating a new Seurat object containing only the cells from the cluster of interest. PCA, clustering, UMAP, and marker gene identification analysis were repeated, as described above.

#### Cell-cell interaction analysis

To infer cell-cell interactions in the tumor stroma we employed CellChat v2 (*43*). The majority of the functions were used according to the standard CellChat workflow for the analysis of single datasets. We used the whole CellChat database for the mouse, including all available interactions. Only genes expressed in at least 1% of total cells were used to infer cell-cell communication. The function identifyOverExpressedGenes() was used with the following parameter settings: thresh.p = 0.05, thresh.fc = 0.2, thresh.pc = 0.1 and the population.size parameter in computeCommunProb() was set to FALSE due to the prior FACS sorting of the cells. Various functions included in the CellChat package were used for the visualization of the results.

#### Trajectory analysis

To perform trajectory analysis on the tumor-associated pericytes, we used monocle3 (v1.3.7) (*49*). Initially, we created a *CellDataSet* object out of the raw expression matrix and the gene annotation, with the new_cell_data_set() command. Next, we performed the main steps of the standard monocle3 workflow, utilizing the preprocess_cds(), reduce_dimension(), cluster_cells() and learn_graph() commands. To identify a root cell for the trajectory, we applied the function get_earliest_principal_node on the ecmPCs subcategory, which we then passed as input to the function order_cells(). Finally plot_cells() was used to visualize the resulting pseudotime trajectory.

#### Human data

The scRNA-seq human datasets that we utilized in this study were available in the NCBI Gene Expression Omnibus (GEO) database under the following GEO accession numbers: GSE144735, GSE132465 (*36*), GSE178341 (*35*), GSE200997 (*38*). The dataset in (*37*) was available as supplementary files (processed gene expression data and metadata) in the original publication. We first processed each dataset separately. In all cases except for Khaliq et al, we used the authors’ cell-type available annotation to only include the stromal cells (Endothelial, Fibroblasts and Pericytes) in the downstream analysis. We then performed our own quality control analysis for each dataset (or cohort in the case of Lee et al) based on the automatic computation of thresholds for the same metrics, as mentioned above. In more detail, we excluded top and bottom outliers for each metric based on the 95% and 5% quantiles of each corresponding distribution. We then merged all datasets including only the intersection of features and then filtered out those expressed in less than 1% of total cells. In the final object the cell number per study was: Pelka et al – 10937, Lee et al – 9653, Qi et al – 7464, Khaliq et al – 474. Downstream analysis was done as described above. To reduce the batch effect across the different datasets, we utilized the command RunHarmony from the Harmony (v1.2.3) package, with the parameters theta equal to 2 and sigma equal to 0.1. Finally, we manually annotated the resulting cell clusters based on top Marker genes. Subclustering of human pericytes was performed as mentioned in the previous sections. We manually annotated the resulting clusters based on the expression of top gene markers and the expression of specific pericyte signatures derived from our analysis in the mouse. We computed signatures by using Ucell (https://github.com/carmonalab/UCell).

## Statistics

Data are presented as mean ± standard deviation (SD). Statistical analyses were performed using GraphPad Prism 10 software. Differences between two groups were analyzed using Student’s t-test. For multiple group comparisons, one-way ANOVA test was used. Welch’s correction was used for samples that showed unequal variance. Differences were considered statistically significant at p ≤ 0.05.

## Data availability

All in-house scRNA-seq data have been deposited in the Gene Expression Omnibus (GEO) database under the accession numbers GSE296872 and GSE296873.

## Acknowledgments

We would like to thank Sofia Grammenoudi for assistance in FACS sorting experiments. We would also like to thank Fleming’s Animal House and Flow Cytometry facilities. This work was funded by Worldwide Cancer Research (Grant No: 22-0126, to VK), and the project Fib3R (Grant No: 3001, to VK) funded by the Hellenic Foundation of Research and Innovation (H.F.R.I.) under the “2^nd^ Call for H.F.R.I. Research Projects to support Faculty members and Researchers”. We acknowledge support of this work by project SingleOut (HFRI-FM17C3- 3780) funded by the Hellenic Foundation for Research and Innovation (H.F.R.I.) under the “1st Call for H.F.R.I. Research Projects to support Faculty members and Researchers and the procurement of high-cost research equipment”. We also acknowledge support by the Research Infrastructure projects InfrafrontierGR (MIS 5002802) and pMedGR (MIS 5002802) funded by the Operational Programme ‘Competitiveness, Entrepreneurship and Innovation’ (NSRF 2014- 2020) and co-financed by Greece and the European Union (European Regional Development Fund), as well as by project MIS 6004752 funded by the Regional Operational Programme ‘ATTICA’ (NSRF 2021-2027) and co-financed by Greece and the European Union (European Regional Development Fund).

## Author contributions

N.C. and V.K. designed the study; N.C. V.Z.A, C.P., and A.M. performed experiments; A.S. performed the bioinformatics analysis; C.N. and V.K. supervised the bioinformatic analysis; N.C., A.S., and V.K. interpreted the experimental results, wrote the manuscript, and prepared the figures; V.K. supervised the study; All authors critically revised and approved the final manuscript.

